# Emergent Simplicity in Microbial Community Assembly

**DOI:** 10.1101/205831

**Authors:** Joshua E. Goldford, Nanxi Lu, Djordje Bajic, Sylvie Estrela, Mikhail Tikhonov, Alicia Sanchez-Gorostiaga, Daniel Segrè, Pankaj Mehta, Alvaro Sanchez

## Abstract

Microbes assemble into complex, dynamic, and species-rich communities that play critical roles in human health and in the environment. The complexity of natural environments and the large number of niches present in most habitats are often invoked to explain the maintenance of microbial diversity in the presence of competitive exclusion. Here we show that soil and plant-associated microbiota, cultivated *ex situ* in minimal synthetic environments with a single supplied source of carbon, universally re-assemble into large and dynamically stable communities with strikingly predictable coarse-grained taxonomic and functional compositions. We find that generic, non-specific metabolic cross-feeding leads to the assembly of dense facilitation networks that enable the coexistence of multiple competitors for the supplied carbon source. The inclusion of universal and non-specific cross-feeding in ecological consumer-resource models is sufficient to explain our observations, and predicts a simple determinism in community structure, a property reflected in our experiments.

## Introduction

The observations that large numbers of microbial species coexist in diverse habitats, ranging from the human gut (*1*) to marine ecosystems (*2*), have renewed interest in understanding the principles that govern the assembly of large microbial communities. Competition amongst microbes for limiting environmental resources is believed to be an important factor shaping community structure. For example, microbes with low competitive ability in local habitats, such as the different human body sites (*3*), are selected against during community assembly in a process known as environmental filtering (*4*). Ecological theory suggests that if species interactions are mediated by competition for resources, then the number of surviving species is bounded by the number of resources in the environment (*5*). In an extreme case, the competitive exclusion principle (*6, 7*) suggests that species whose growth is limited by the same single environmental resource cannot coexist in that environment without additional compensatory mechanisms (*5*). Indeed, mathematical models of large ecosystems of competing species suggest that the ensuing communities are intrinsically unstable, unless specific conditions, like non-transitive competition effects, are met (*8, 9*). For this reason, the complexity of natural environments, which is characterized by either the supply of large numbers of different resources (*10*), spatial and temporal structure (*11*), or the presence of biotic interactions such as phage predation (*12*), is often thought to underlie the maintenance of microbial diversity in large ecosystems (*5*).

Although most of the work on community assembly has focused on competition (*13, 14*), facilitation may also be an important force that structures large ecological communities (*15–18*). Facilitation may be particularly important amongst microbes, whose metabolism involves the secretion of byproducts that increase environmental complexity, which potentially creates new niches leading to syntrophy (*19, 20*). Several examples of facilitation-mediated coexistence on a single limiting nutrient are known (*21–23*), including cases where syntrophic interactions emerge *de novo* in long-term evolutionary experiments (*24–26*), making it reasonable to hypothesize that these interactions are common within natural microbial communities.

To determine whether facilitation-mediated coexistence on a single limiting nutrient is indeed a universal phenomenon, we developed a high-throughput *ex situ* cultivation protocol to monitor the spontaneous assembly of ecologically stable microbial communities derived from natural habitats. By precisely manipulating the chemical composition of synthetic growth media, we were able to systematically investigate the interplay between competition and facilitation during environmental filtering, and gain insight into the rules of microbial community assembly in large communities.

### Stable microbial communities on a single limiting resource

We devised a top-down approach to monitor community assembly in high-throughput, in which we forced naturally existing communities to re-assemble in synthetic (M9) minimal media containing a single externally-supplied source of carbon (Methods) as well as single sources of all of the necessary salts and chemical elements required for microbial life (fig 1a). Intact microbiota suspensions were extracted from diverse natural ecosystems, such as various soils and plant leaf surfaces (Methods). Suspensions of microbiota from these environments were highly diverse and taxonomically rich (fig. S1), ranging between 110 and 1290 exact sequence variants (ESV). We first inoculated 12 of these suspensions of microbiota into fresh minimal media with glucose as the only added carbon source, and allowed the cultures to grow at 30°C in static broth. We then passaged the mixed cultures in fresh media every 48 hours with a fixed dilution factor of *D* = 8×10^−3^ for a total of 12 transfers (~84 generations). At the end of each growth cycle, we assayed the community composition using 16S rRNA amplicon sequencing (fig. 1a, Methods). High resolution sequence denoising allowed us to identify ESVs, which revealed community structure at single nucleotide resolution (*27*).

Most communities stabilized after ~60 generations, reaching stable population equilibria in nearly all cases (fig. 1b, S2). For all of the 12 initial ecosystems, we observed large multispecies communities after stabilization that ranged from 4 to 17 ESVs at a sequencing depth of 10,000 reads (fig. S3-4, see Methods). We confirmed the taxonomic assignments generated from amplicon sequencing by culture-dependent methods, including 87 the isolation and phenotypic characterization of all dominant genera within a representative community (fig. S5).

**Figure 1:**
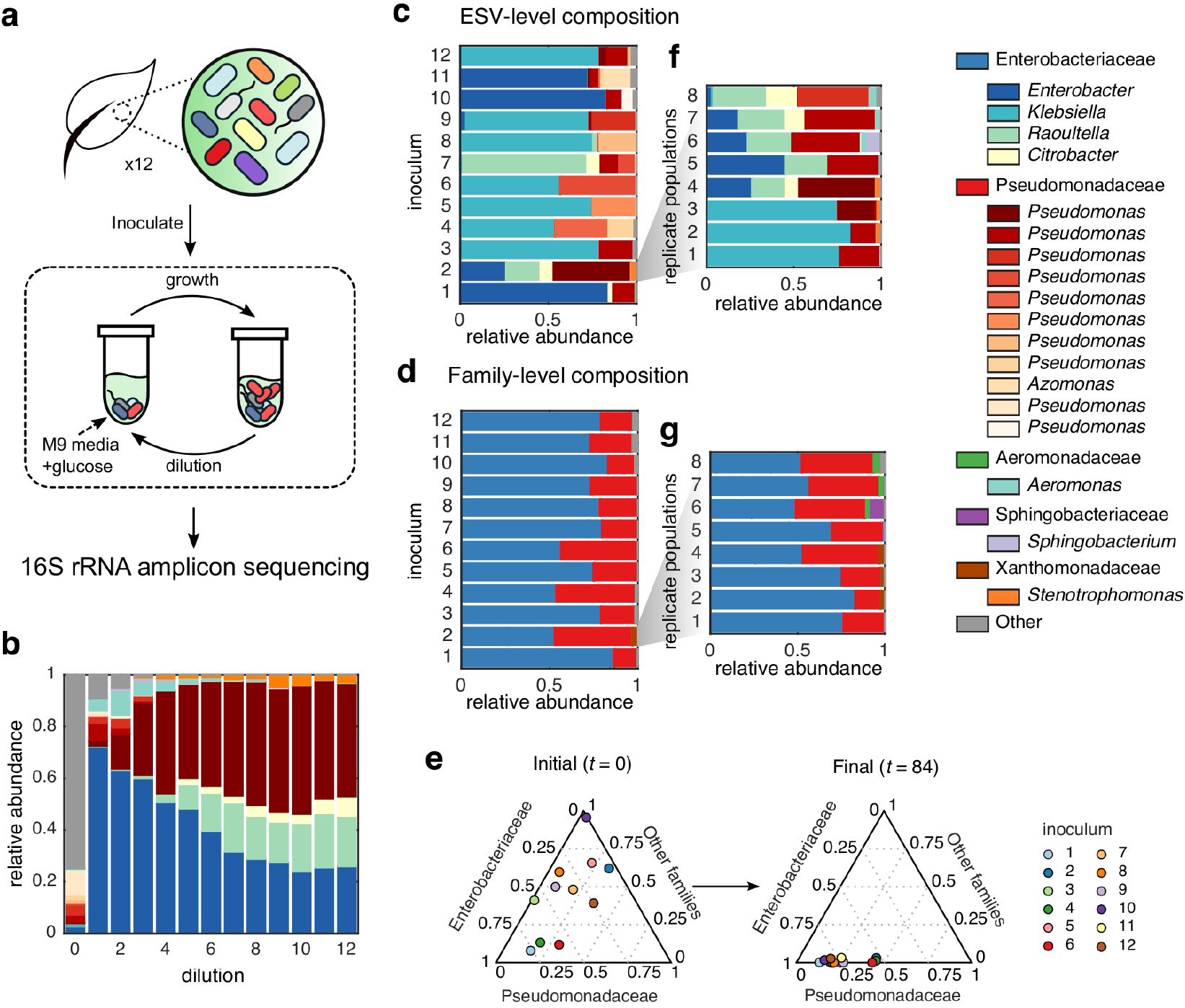
Top down assembly of bacterial consortia. (a) Experimental scheme: large ensembles of taxa were obtained from 12 leaf and soil samples, and used as inocula in passaged-batch cultures containing synthetic media supplemented with glucose as the sole carbon source. After each transfer, 16S rRNA amplicon sequencing was used to assay bacterial community structure. (b) The community structure of a representative community (from inoculum 2) after every dilution cycle (~7 generations), revealing a 5-member consortia from the *Enterobacter, Raoultella, Citrobacter, Pseudomonas* and *Stenotrophomonas* genera. The community composition after 84 generations is shown at the exact sequence variant (ESV) level (c) or the family taxonomic level, converging to characteristic fractions of Enterobacteriaceae and Pseudomonadaceae (d) (Monte Carlo permutation test, *P* < 10^−4^). (e) Simplex representation of family-level taxonomy before (*t* = 0) and after (*t* = 84) passaging experiment. (f-g) Experiments were repeated with 8 replicates from a single source (inocula 2), and communities converged to very similar family level distributions (g), but displayed characteristic variability at the genus and species level (f).

### Convergence of bacterial community structure at the family taxonomic level

High-throughput isolation and stabilization of microbial consortia allowed us to explore the rules governing the assembly of bacterial communities in well-controlled synthetic environments. At the species (ESV) level of taxonomic resolution, the 12 natural communities assembled into highly variable compositions (fig. 1c). However, when we grouped ESVs by higher taxonomic ranks we found that all 12 stabilized communities, with very diverse environmental origins, converged into highly similar family-level community structures dominated by Enterobacteriaceae and Pseudomonadaceae (fig. 1d). In other words, the same family-level composition arose in all communities despite their very different starting points, indicating the existence of a dynamic attractor at the family taxonomic level (Monte Carlo permutation test, *P* < 10^−4^, fig. 1d-e), which was not observed when communities were examined at the genus or species levels (Monte Carlo permutation test, *P* = 0.12, Methods).

To better understand the origin of the taxonomic variability observed below the family-level, eight replicate communities were started from each one of the 12 starting microbiome suspensions (inocula), and propagated in minimal media with glucose as in the previous experiment. Given that the replicate communities were assembled in identical habitats and were inoculated from the same pool of species, any observed variability in community composition across replicates would suggest that random colonization from the regional pool and microbe-microbe interactions are sufficient to generate alternative species-level community assembly. Indeed, for most of the inocula (nine out of twelve), replicate communities assembled into alternative stable ESV-level compositions, while still converging to the same family-level attractor described in fig. 1e (see also fig. S6). One representative example is shown in fig. 1f-g; all eight replicates from the same starting inoculum assemble into strongly similar family-level structures, which are quantitatively consistent with those found before (fig. 1d). However, different replicates contain alternative Pseudomonadaceae ESVs, and the Enterobacteriaceae fraction is constituted by either an ESV from the *Klebsiella* genus, or a guild consisting of variable subcompositions of *Enterobacter, Raoultella*, and/or *Citrobacter* as the dominant taxa. The remaining (three out of twelve) inocula reached highly convergent community compositions, with all replicates exhibiting strongly similar population dynamics and population structures at all levels of taxonomic resolution (fig. S7). The reproducibility in population dynamics between replicate communities ensures that experimental error is not the main source of variability in community composition.

Despite the observed species level variation in community structure, the existence of family-level attractors suggests the existence of fundamental rules governing community assembly. Recent work on natural communities has consistently found that environmental filtering selects for convergent function across similar habitats, while at the same time allowing for taxonomic variability within each functional class (*28, 29*). In our assembled communities in glucose media, fixed proportions of Enterobacteriaceae and Pseudomonadaceae may have emerged due to a competitive advantage, given the well-known glucose uptake capabilities of the phosphotransferase system in Enterobacteriaceae and ABC transporters in Pseudomonadaceae (*30*). This suggests that the observed family-level attractor may change if we assemble communities adding a different carbon source to our synthetic media. To determine the effect of the externally provided carbon source on environmental filtering, we repeated the community assembly experiments with eight replicates of all 12 natural communities, using two alternative single-carbon sources, citrate or leucine, instead of glucose. Consistent with previous experiments on glucose minimal media, communities assembled in citrate or leucine contained large numbers of species: communities stabilized on leucine contained 6-22 ESVs, while communities stabilized on citrate contained 4-22 ESVs at a sequencing depth of 10,000 reads. As was the case for glucose, replicate communities assembled on citrate and leucine also differed widely in their ESV-level compositions, while converging to carbon source-specific family-level attractors (fig. 2a, S8-9). Family-level community similarity (Renkonen similarity) was on average higher between communities passaged on the same carbon source (median: 0.88) than between communities passaged from the same environmental sample (median 0.77, one-tailed Kolmogorov-Smirnov test, *P* < 10^−5^; fig. S10). Communities stabilized in citrate media were composed of a significantly lower fraction of Enterobacteriaceae (Mann-Whitney U test, *P* < 10^−5^), and displayed an enrichment of Flavobacteriaceae relative to communities grown on glucose (Mann-Whitney U test, *P* < 10^−5^), while communities stabilized in leucine media had no growth of Enterobacteriaceae and an enrichment for Comamonadaceae relative to communities grown on glucose (Mann-Whitney U test, *P* < 10^−5^) or citrate (Mann-Whitney U test, *P* < 10^−5^).

These results suggest that the supplied source of carbon exerts a strong environmental filter on community assembly. To quantify this effect, we used a machine learning approach and trained a support vector machine (SVM) to predict the identity of the supplied carbon source from the family-level community composition. We obtained a cross-validation accuracy of 97.3% (fig. 2b; Methods). Importantly, we found that considering the tails of the family-level distribution (as opposed to just the two dominant taxa) increases the predictive accuracy (fig. 2b), which indicates that carbon source-mediated environmental filtering extends to the entire family-level distribution, including the more rarefied members.

**Figure 2:**
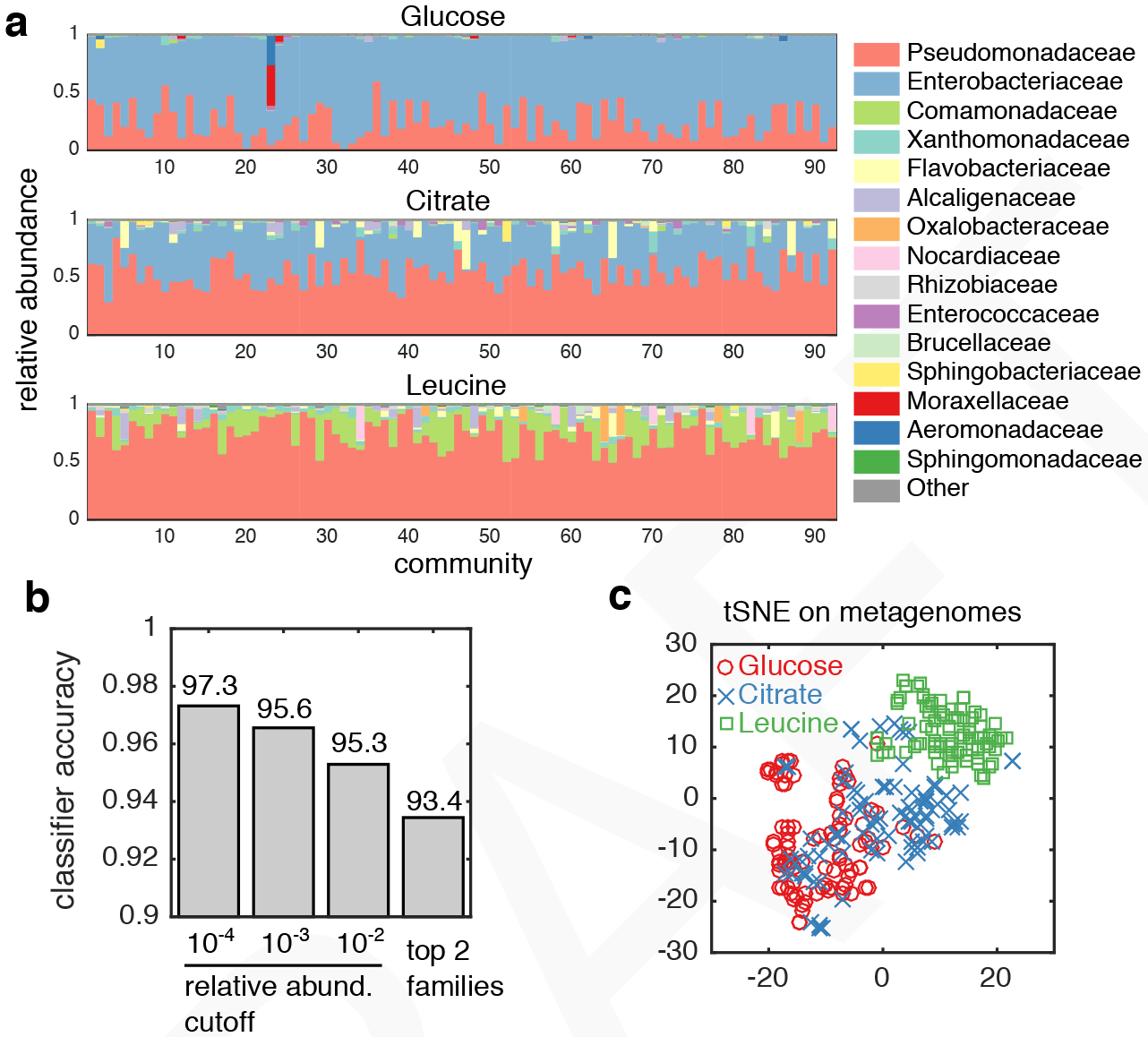
Family-level and metagenomic attractors are associated with different carbon sources. (a) Family-level community compositions are shown for all replicates across 12 inocula grown on either glucose, citrate or leucine as the limiting carbon source. (b) A support vector machine (see Methods) was trained to classify the carbon source from the family-level community structure. Low abundant taxa were filtered using a predefined cutoff (*x*-axis) before training and performing 10-fold cross validation (averaged 10 times). Classification accuracy with only Enterobacteriaceae and Pseudomonadaceae resulted in a model with ~93% accuracy (right bar), while retaining low abundant taxa (relative abundance cutoff of 10^−4^) yielded a classification accuracy of ~97% (left most bar). (c) Metagenomes were inferred using PICRUSt (*50*), and dimensionally reduced using tSNE, revealing that carbon sources are strongly associated with the predicted functional capacity of each community.

Substantial environmental filtering is thus exerted by the supplied carbon source, and it leads to quantitative rules of assembly. Rather than selecting for the most fit single species, environmental filtering leads to communities that contain fixed fractions of multiple coexisting families that are determined by the carbon source in a strong and predictable manner (fig. S10). This suggests the existence of underlying mechanisms of taxonomic selection by function. Consistent with this idea, we find that the inferred community metagenomes assembled on each type of carbon source exhibit substantial clustering by the supplied carbon source (fig. 2c), and are enriched in pathways for its metabolism (fig. S16). Interestingly, when we spread the stabilized communities on agarose plates, we routinely found multiple identifiable colony morphologies per plate, evidencing that multiple taxa within each community are able to grow independently on (and thus compete for) the single supplied carbon source. This suggests that the genes and pathways that confer each community with the ability to metabolize the single supplied resource are distributed among multiple taxa in the community, rather than being present only in the best competitor species.

### Widespread metabolic facilitation stabilizes competition and promotes coexistence

Classic consumer-resource models indicate that when multiple species compete for a single externally supplied growth-limiting resource, the only possible outcome is competitive exclusion unless specific circumstances apply (*5, 8–12*). However, this situation does not adequately reflect microbes, whose ability to engineer their own environments is well documented both in the lab (*20, 25, 31, 32*) and in nature (*33, 34*). Thus, we hypothesized that the observed coexistence of competitor species in our experiments may be attributed to the generic tendency of microbes to secrete metabolic byproducts into the environment, which could then be reutilized by other community members.

To determine the plausibility of niche creation mediated by metabolic byproducts, we analyzed one representative glucose community in more depth. We isolated members of the four most abundant genera in this community (*Pseudomonas, Rauoultella, Citrobacter* and *Enterobacter*), which together represented ~ 97% of the total population in that community (fig 3a). These isolates had different colony morphologies and were also phenotypically distinct (fig. S5). All isolates were able to form colonies in glucose agarose plates and all grew independently in glucose as the only carbon source, which indicates that each isolate can compete for the single supplied resource. To test the potential for cross-feeding interactions in this community, we grew monocultures of the four isolates for 48 hours in synthetic M9 media containing glucose as the only carbon source (fig. 3b). After 48 hours, the glucose concentration was too low to be detected, indicating that all of the supplied carbon had been consumed and any carbon present in the media originated from metabolic byproducts previously secreted by the cells. To test whether these secretions were enough to support growth of the other species in that community, we filtered the leftover media to remove cells, and added it to fresh M9 media as the only source of carbon (fig. 3b). We found that all isolates were able to grow on every other isolate’s secretions (e.g. fig. 3c), forming a fully connected facilitation network (fig. 3d). Growth on the secretions of other community members was strong, often including multiple diauxic shifts (fig. S11), and the amount of growth on secretions was comparable to that on glucose (fig. S12), suggesting the pool of secreted byproducts are diverse and abundant in this representative community.

To find out if growth on metabolic byproducts is frequent among our communities, we thawed 95 glucose-stabilized communities (7-8 replicates from 12 initial environmental habitats) and grew them again on glucose as the only carbon source for an extra 48 hour cycle. In all 95 communities glucose was completely exhausted after 24 hours of growth (fig. 3e); yet, most communities continued growing after glucose had been depleted (fig. 3e), showing that growth on previously secreted byproducts is widespread. Moreover, community growth on the secreted byproducts is strong: on average, communities produce ~25% as much biomass on the secretions alone as they did over the first 24 hours when glucose was present (fig. 3f). Growth after glucose depletion is on average ~5-fold higher than the proportion of measured dead cells in representative communities (fig. S13), supporting the hypothesis that metabolic byproduct secretion (rather than cell lysis) is the dominant source of the observed cross-feeding. Other mechanisms may also operate together with facilitation in specific communities to support high levels of biodiversity (*9, 35–38*). Surprisingly, we found that multi-species communities still form in the absence of spatial structure, and we did not observe effects from temporal competitive niches in our experiments (fig. S14-15). Although beyond the scope of this work, efforts to elucidate the roles of other mechanisms that may stabilize competition, like phage predation (*12*) or non-transitive competition networks (*39*), will more fully characterize the landscape of interactions in these microcosms.

**Figure 3:**
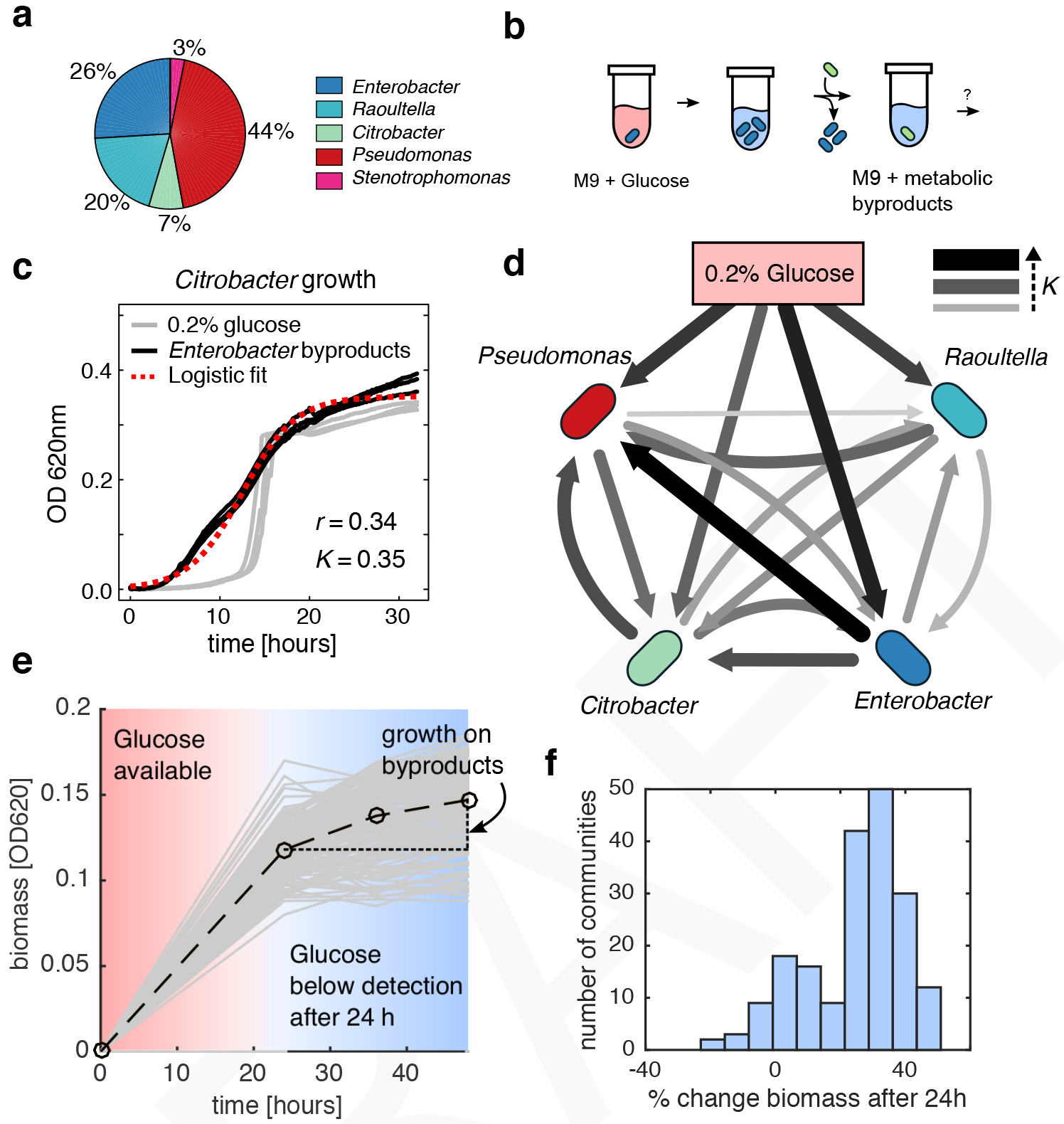
Non-specific metabolic facilitation stabilizes competition for the supplied resource. (a) The major taxa in a representative community from inoculum 2 were isolated, grown under conditions with minimal media (M9) and glucose, and the metabolic byproducts were used as the sole carbon source in growth media for other isolates. (b) Experimental set-up: isolates were grown in minimal media with glucose for 48 hours, and cells were filtered out from the suspension. The suspension of byproducts was mixed 1:1 with 2X M9 media and used as the growth media for other isolates (see also Methods). (c) An example growth curve for *Citrobacter* growing either with M9 supplemented with 0.2 % glucose (grey line) or the metabolic byproducts from *Enterobacter* (black line). (d) All isolates were grown on every other isolate’s metabolic byproducts, and logistic models were used to fit growth curves. We plotted the fitted growth parameters (carrying capacity) as edges on a directed graph, where the edges encode the carrying capacity of the target node isolate when grown using the secreted byproducts from the source node isolate. Edges from the top node encodes the carrying capacity on 0.2 % glucose, which is comparable in edge width/color to several of the other interactions. (e-f) All communities stabilized on glucose were grown in glucose-supplemented M9 media, and optical densities at 620 nm were measured, showing that after glucose was depleted (~24 hours), communities on average grew an additional 25%.

### Including cross-feeding in a generic consumer resource model recapitulates experimental observations

The above experiments suggest that competition for the single supplied limiting nutrient may be stabilized by non-specific metabolic facilitation, leading to coexistence. To test whether this feature alone could promote widespread coexistence, we simulated a community assembly process on a single supplied carbon source using a version of the classic MacArthur consumer resource model (CRM) (*40*), which was modified to include non-specific cross-feeding interactions. Cross-feeding was modeled through a stoichiometric matrix that encodes the proportion of a consumed resource that is secreted back into the environment as a metabolic byproduct (Supporting Information). Setting this matrix to zero results in no byproducts being secreted, and recovers the classic results for the CRM in a minimal environment with one resource: the species with highest consumption rate of the limiting nutrient competitively excludes all others (fig. 4a, inset). However, when we drew the stoichiometric matrix from a uniform distribution (while ensuring energy conservation), and initialized simulations with hundreds of “species” (each defined by randomly generated rates of uptake of each resource) coexistence was routinely observed (fig. 4a). All of the coexisting “species” in this simulation were generalists, capable of growing independently on the single supplied resource as well as on each other species’ secretions.

Our experiments have shown that the family-level community composition is strongly influenced by the nature of the limiting nutrient, which may be attributed to the metabolic capabilities associated with each family. We modeled this scenario by developing a procedure that sampled consumer coefficients from four metabolic “families”, ensuring that consumers from the same family were metabolically similar (see Supporting Information). We randomly sampled a set of 100 consumer vectors (or “species”) from four families, then simulated growth on 20 random subsets of 50 species on one of three resources (labeled here as *A, B* or *C*). As in our experimental data (fig. 2a), simulated communities converged to similar family-level structures (fig. 4c), despite displaying variations at the species level (fig. 4b). We confirmed the correspondence between family-level convergence and functional convergence by computing the community-wide metabolic capacity per simulation, resulting in a predicted community-wide resource uptake rate for each resource (Supporting Information). Communities grown on the same resource converged to similar uptake capacities with an enhanced ability to consume the limiting nutrient (fig. 4d). Importantly, this functional convergence is exhibited even when consumers are drawn from uniform distributions, with no enforced family-level consumer structure, suggesting that the emergence of functional structure at the community level is a universal feature of consumer resource models (fig. S16). Strikingly, the composition of surviving species within each metabolic family frequently displayed “guilds” of species capable of supporting the stable growth of rare (<1% relative abundance) taxa, rather than a single representative from each family (fig. 4e), similar to our experimental data (fig. 1c,e). Our model suggests that guilds of species are stabilized by a dense facilitation network (fig. 4f), consistent with observations of widespread metabolic facilitation in experiments (fig. 3d). Thus, we find that simulations of community dynamics with randomly generated metabolisms and resource uptake capabilities capture a wide range of qualitative observations found in our experiments.

**Figure 4:**
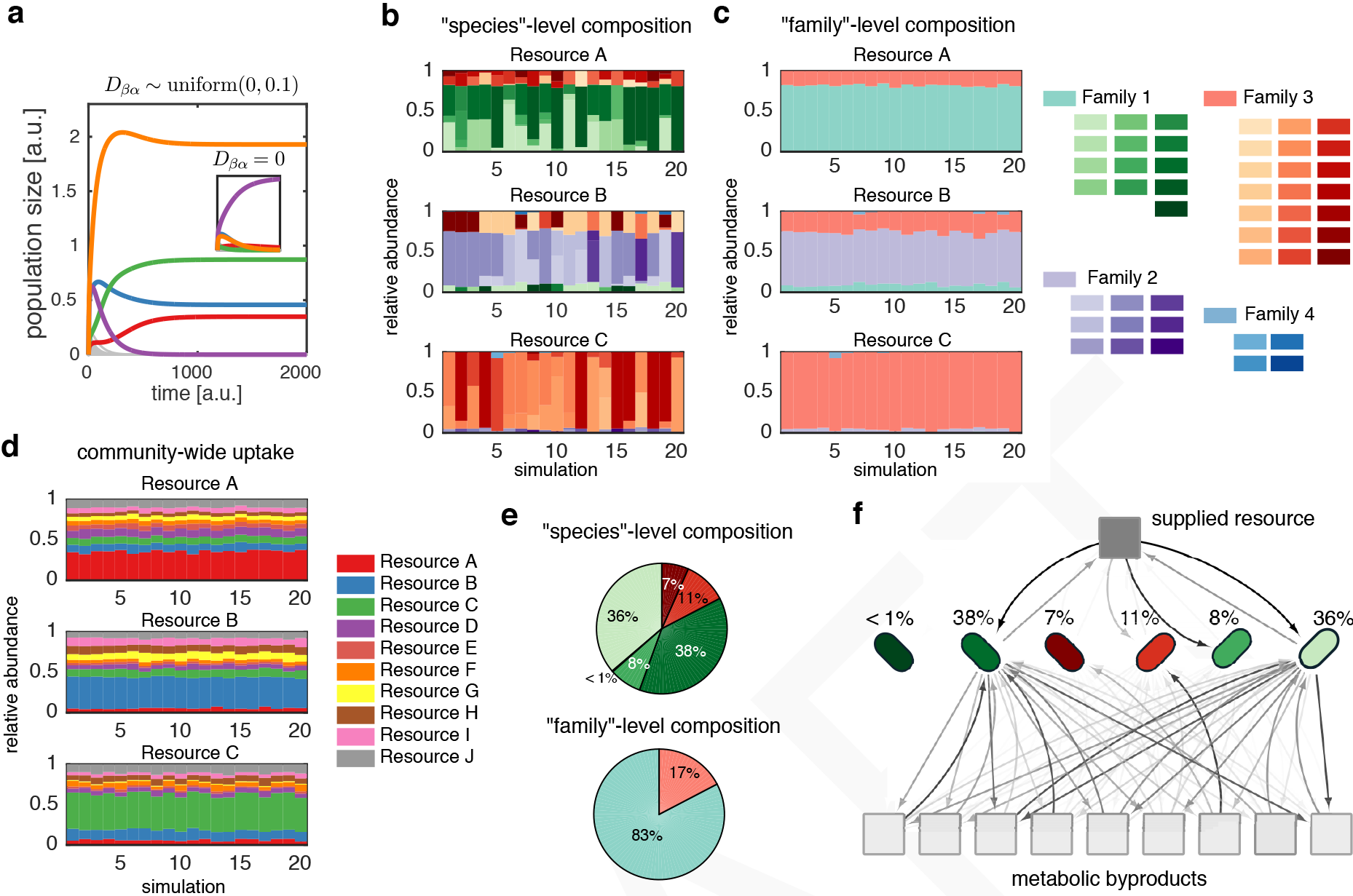
A simple extension of classic ecological models recapitulates several experimental observations. MacArthur’s consumer resource model was extended to include 10 byproduct secretions along with consumption of a single primary limiting nutrient (see Supporting information), controlled by a global stoichiometric matrix *D*_βα_, which encodes the proportion of the consumed resource α that is transformed to resource β and secreted back into the environment. Consumer coefficients were sampled from 4 characteristic prior distributions, representing four “families” of similar consumption vectors. (a) Simulations using a randomly sampled global stoichiometric matrix generically resulted in coexistence of multiple competitors, while setting this matrix zero eliminated coexistence (a, inset). Random ecosystems often converged to very similar family-level structures (c), despite variation in the species-level structure (b). The family-level attractor changed when providing a different resource to the same community (b-c, subplots). (d) Total resource uptake capacity of the community was computed (Supporting Information), analogous to the inferred metagenome (see fig. 2d), and is, like the family-level structure, highly associated with the supplied resource. (e) Communities that formed did not simply consist of single representatives from each family, but often consisted of guilds of several species within each family, similar to experimental data. (f) The topology of the flux distribution shows that surviving communities all compete for the primary nutrient and competition is stabilized by differential consumption of secreted byproducts. The darkness of the arrows corresponds to the magnitude of flux.

## Discussion

The theory and experiments described above allow us to evaluate how the interplay between competition and facilitation governs microbial community assembly. Our results indicate that assembly of large communities is to be expected even in simple nutrient limited environments. We find substantial evidence that generic, non-specific facilitation stabilizes competition, enabling the coexistence of multiple taxa that compete for a single supplied limiting resource. Overall, our results indicate that resource-mediated facilitation and competition are not easily separable in microbial communities. The theoretical and computational framework for modeling community assembly is largely based on networks of pairwise competitive or facilitative interactions (*41–44*). Although recent work has called attention upon higher-order interactions (*8*, *39*), the extent to which they affect microbial community assembly remains poorly understood. Our results suggest that collective interactions, such as those naturally arising when multiple species collectively transform their environment by consuming and secreting resources, may generally play an important and underappreciated role in the dynamics and assembly of large microbial ecosystems. Future work should shed light on the implications and full extent of collective interactions in microbial ecosystems.

The mechanisms of microbe-microbe interactions are not limited to competition for resources, and they can include a wide range of direct and indirect physical, chemical and biological processes, from detoxification to predation or killing by contact (*45–48*). The range of processes that may simultaneously operate in any one community, and the non-linear nature of their effects, introduce significant complications for the bottom-up prediction of community assembly outcomes at the species level, particularly for large communities with large number of species (*17*). As an illustration of these challenges, we found that even in replicate habitats that are inoculated from the same starting natural microbiota, communities often assemble into different alternative states at the species level (fig. 1f). Encouragingly, however, we find that the opposite is true at higher orders of taxonomic description. When we group taxa by family, reassembly from widely different initial communities exhibits an emergent simplicity: while the specific mechanisms of interaction between individual strains in each community (or even the starting inoculum) may vary widely across communities, coarse-grained community assembly is deterministic and predictable, and functionally induced by the type of carbon source available in the media.

The same quantitative assembly rules for large communities also emerge in generic consumer-resource simulations when they are initialized with species with random metabolisms, agnostic to specific mechanisms of species interactions in each community. This suggests that this emergent simplicity at higher levels of organization is a generic property of large, diverse ecosystems. Thus, we hypothesize that many of the patterns observed in recent large-scale microbiome studies - for example, the convergence of the metagenome and phylum-level structure in systems as diverse as the human gut (*1, 49*), plant foliage (*28*) or the oceans (*2*) - may reflect universal properties of large self-sustained microbial communities.

## Acknowledgments

The funding for this work partly results from a Scialog Program sponsored jointly by Research Corporation for Science Advancement and the Gordon and Betty Moore Foundation through grants to Yale University and Boston University by the Research Corporation and by the Simons Foundation. This work was also supported by NIH NIGMS grant 1R35GM119461, a Simons Investigator award in the Mathematical Modeling of Living Systems (MMLS). DS and JEG additionally acknowledge funding from the Defense Advanced Research Projects Agency (Purchase Request No. HR0011515303, Contract No. HR0011-15-C-0091), the U.S. Department of Energy (DE-SC0012627), the NIH (T32GM100842, 5R01DE024468, R01GM121950 and Sub_P30DK036836_P&F), the National Science Foundation (1457695), the Human Frontiers Science Program (RGP0020/2016), and the Boston University Interdisciplinary Biomedical Research Office. We wish to express our gratitude to the Goodman laboratory at Yale, and to the Brucker laboratory at the Rowland Institute for their technical help on the early phases of this project, and to members of the Sanchez, Mehta and Segrè groups for helpful discussions.

## Methods

### Isolating microbial communities from natural ecosystems

Leaf or soil samples (~1 g) were collected from natural environments using sterile tweezers and placed in 15 mL falcon tubes. In the lab, 10 mL of 5% NaCl buffer was added to each sample and allowed to incubate for ~48 hours at room temperature. 40% glycerol stock solutions were prepared from aqueous sample suspensions and frozen at −80 °C for storage.

### Preparation of 96-well media plates

All media contained 0.07 C-mole/L of carbon source (glucose, citrate or leucine) and was sterile-filtered with a 0.22 μm filter (Millipore). Stock solutions of carbon sources were stored at 4°C for no more than 1 month. M9 media was prepared from concentrated stocks of M9 salts (without MgSO_4_ or CaCl_2_) and stock solutions of MgSO_4_ and CaCl_2_. 500 μL cultures containing 450 μL of sample and 50 μL stock carbon source were grown in 96 deep-well plates (VWR). For the first two cell passages, cycloheximide was added to the media at a concentration of 200 μg/mL to inhibit eukaryotic growth.

### Passaging microbial populations

Starting inocula were obtained directly from the initial buffered solution of microbiota by inoculating 4 μL into 500 μL culture media. For each sample, 4 μL of the culture medium was dispensed into all 60 wells of the fresh media plate. Cultures were allowed to grow for 48 hours at 30 °C in static broth, then each culture was triturated 10 times to ensure communities were homogenized before passaging. Passaging was performed by taking 4 μL from each culture to use as inocula in 500 μL of fresh media, and cells were allowed to grow again. Cultures were passaged 12 times (~84 generations). Optical density (OD_620_) was used to measure biomass in cultures after the 48-hour growth cycle. Samples to be sequenced were collected and stored by spinning down in a micro-centrifuge for 10 min at 14,000 RPM at room temperature. Cell pellets were stored at −20 °C.

### Fermentation assays and isolation of strains

Four bacterial strains from a representative community stabilized in glucose were isolated and identified taxonomically. The community was plated onto 0.5% agarose Petri-dishes containing M9 supplemented with 0.2% glucose and were allowed to grow for 48 hours at 30°C. Single colonies were then picked from these plates according to their colony morphologies, re-streaked on fresh agarose plates and grown for another 48 hours at 30°C. Single colonies from each isolate grown for 48 hours at 30°C in liquid M9 supplemented with 0.2% glucose were finally stored at −80 °C in 40% glycerol. Isolates were also identified according to their differential ability to ferment the following 16 carbohydrates: adonitol, arabinose, cellobiose, dextrose, dulcitol, fructose, inositol, lactose, mannitol, mannose, melibiose, raffinose, rhamnose, salicin, sucrose, and xylose (fig S5 a-b). Fermentation ability was assessed using a phenol red broth base with an added carbohydrate at a final concentration of 1% w/v, except for cellobiose (0.25%) due to its low solubility. Each isolate was grown on an agarose plate, and a single colony was picked and re-suspended into 100 μL 1x PBS. 2 μL of each isolate was inoculated into 50 μL of Phenol red broth + carbon source (in a 384 well-plate, Corning). Spectrophotometric measurements of phenol red (OD_450_ and OD_551_) were measured after 0h, 12, 16, and 19 hours of incubation. Clustering of O.D. profiles after 19 hours revealed 4 distinct phenotypic profiles, consistent with morphologies (fig. S5c). Taxonomic assignments of isolates were verified using full-length 16S rRNA sequencing of DNA extracted from single colonies grown on agarose plates (GENEWIZ), using the online RDP classifier (*51*).

### Metabolic facilitation assay and measurement of glucose depletion

To determine whether microbial cross-feeding is a potential mechanism that enables coexistence, four isolates from a single representative community were inoculated in 5 mL of M9 media with 0.2% glucose, then incubated for 48 hours at 30 °C (fig. 3a). Cells were then separated from the spent media (SM) using the following procedure: cells were centrifuged at 3000 rpm for 10 min, and SM was filter-sterilized and stored at 4 °C. Cells were re-suspended in the same volume of PBS, and washed two times times by centrifugation (3000rpm, 10min). Cells were diluted to an OD_620_ of 0.24 prior to inoculation. There was no detectable glucose remaining in any SM as measured using the Glucose GO Assay Kit (Sigma), with the exception of the SM from *Pseudomonas*, which was adequately controlled for (see main text). SM was then mixed 1:1 with fresh 2X M9 media with no carbon source. Each isolate was inoculated in each isolate’s SM-based M9 in triplicate at 1% v/v in a 384 well plate (Corning). The plate was incubated in a standard plate reader (Thermo 498 Scientific), and OD_620_ was measured every 10 min at 30 °C.

We sought to determine whether glucose-stabilized communities were able to grow after glucose depletion, which would suggest that biomass accumulation is attributed to consumption of metabolic byproducts. For this, 95 glucose-stabilized communities were inoculated in a 96 deep-well plate from frozen stock in 500 μL of M9 0.2% glucose. Two initial transfers with 48 hours incubation were performed as previously described (30 °C no shaking). The third transfer was performed in duplicate and with final volume 600 μL. From these two plates, 100 μL samples were taken at 24, 36, 48 and 56 hours. OD_620_ was measured, followed by the measurement of glucose using the Glucose GO Assay Kit (Sigma). Glucose concentrations were inferred using linear regression from the standard curve, although no sample at any time point showed detectable levels.

### Cell death measurements

Samples were obtained at 12-hour intervals to measure the accumulation of biomass and determine the frequency of dead cells. Bacteria stained with the LIVE/DEAD BacLight Bacterial Viability Kit (L-7012, Invitrogen) following manufacturer instructions were spotted on 1% agarose pads. Microscopy was performed on an Eclipse Ti-E microscope (Nikon, Tokyo, Japan), equipped with Perfect Focus System (Nikon), a phase-contrast objective Plan Apochromat 100X/1.40 NA (Nikon), and an ORCA-Flash4.0 V2 Digital CMOS camera (Hamamatsu Photonics, Hamamatsu City, Japan). Red fluorescence of dead cells was recorded with a Texas Red bandpass filter. Images were acquired with MetaMorph software (Molecular Devices, Sunnyvale, CA, USA) and analyzed with Microbe J (*52*).

### DNA extraction, library preparation, sequencing and analysis

Cell pellets were re-suspended and incubated at 37 °C for 30 min in enzymatic lysis buffer (20 mM Tris-HCl, 2mM sodium EDTA, 1.2% Triton X-100) and 20 mg/mL of lysozyme from chicken egg white (Sigma-Aldrich) to lyse the cell walls of Gram-positive bacteria. Following cell lysis, the DNA extractions were performed following the DNeasy 96 protocol for animal tissues (Qiagen). The clean DNA was eluted in 100 μL elution buffer of 10 mM Tris-HCl, 0.5 mM EDTA at pH 9.0. DNA concentration was quantified using Quan-iT PicoGreen dsDNA Assay Kit (Molecular Probes, Inc.) and normalized to 5 ng/μL for subsequent 16S rRNA sequencing. 16S rRNA amplicon library preparation was conducted using a dual-index paired-end approach developed by Kozich et al. (*53*). Briefly, PCR-amplified libraries were prepared using dual-index primers (F515/R806) to generate amplicons spanning the V4 region of the 16S rRNA gene, then pooled and sequenced using the Illumina MiSeq platform. For each sample, a 30-cycle PCR was performed in duplicate in 20 μL reaction volumes using 5 ng of DNA, dual index primers, and AccuPrime Pfx SuperMix (Invitrogen). Thermocycling conditions consisted of a 2-min initial denaturation step at 95 °C, followed by 30 cycles of the following PCR scheme: (a) 20-second denaturation at 95 °C, (b) 15-second annealing at 55 °C, and (c) 5-min extension at 72 °C. PCR was terminated after a 10-min extension step at 72 °C. After pooling amplicons from duplicate reactions, the PCR products were purified and normalized using the SequalPrep PCR cleanup and normalization kit (Invitrogen). Libraries were then pooled and sequenced using Illumina MiSeq v2 reagent kit, which generated 2x250 base pair paired-end reads at the Yale Center for Genome Analysis (YCGA). For shaking control experiments (fig. S15), library preparation and sequencing was performed at SeqMatic (Fremont, CA). Sequencing and library preparation were identical when compared to the procedure described above, except primers targeted the V3-V4 region of 16S rRNA gene.

QIIME 1.9.0 (*54*) was used to demultiplex and remove barcodes, indexes and primers from raw files, producing FASTQ files with for both the forward and reverse reads for each sample. Dada2 v. 1.1.6 (*27*) was used to infer unique sequence variants from each sample, and naïve Bayes was used to assign taxonomy using the SILVA v. 123 database (*51, 55*). Metagenome inference was performed using PICRUSt (*50*). Denoised ESVs were assigned to OTUs using the greengenes database (version 13.5) using the QIIME function pick_closed_reference_otus, with a 97% similarity cutoff. Communities were normalized using the normalize_otus.py function in PICRUSt, and the metagenomes were estimated using the estimate_metagenome.py routine.

### Statistical tests for community convergence

We developed a Monte Carlo permutation test to determine if a group of stabilized communities converged to a consistent community structure at a defined level of taxonomic rank. For each community, we first grouped sequence variants by either genus or family taxonomic rank, then transformed relative genus or family abundances using a centered log-ratio transform (clr) (*3, 56*). We then calculated the variance for each taxonomic label (genus or family) across all samples and summed over all taxa, resulting in a metric we call *h*_rank_, where rank was either genus or family. This metric captures the total variability of the community at pre-defined levels of taxonomic rank. To determine if this variability is lower than what we would expect by chance, we permuted the labels for the sequence variants 10^4^ times and computed 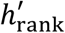 for each random permutation. The probability that a low community-wide variance could be observed by chance 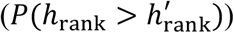 was estimated by computing the fraction of instances when *h*_rank_ exceeded 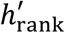.

### Prediction of media carbon source from community structure

To access the predictive quality of the community structure and inferred metagenomes, we trained and evaluated multi-class support vector machine (SVM) models. SVMs were constructed using the MATLAB function fitecoc and evaluated using 10-fold cross validation in fig 2b or leave one out cross-validation in fig. S16. Features used in the in the SVM were either the clr-transformed relative abundances at the family taxonomic level in fig. 2b or the clr-transformed inferred metagenome composition in fig. S16.

### Low abundant growth with no supplied carbon source

Passaging experiments were performed using M9 synthetic media with no additional carbon sources, which resulted in the stabilization of very low abundant microbial communities (fig S4). Growth was often several orders of magnitude lower than growth on either the primary nutrient (fig S4c) or secreted byproducts (fig 3e-f), suggesting that metabolic consumption of secreted byproducts is more likely to contribute to stabilizing competition than consumption of low levels of latent resources in the D.I. water. To determine community richness resulting from growth on the provided resource, we estimated the abundance of 16S amplicon reads deriving from contamination either by cross-well contamination or microbial growth on the low levels of total organic carbon in D.I. water (fig. S4a-b). For each of the 12 initial points, communities propagated for 84 generations with either with M9 and 0.2% glucose, or M9 and no additional carbon source. We plated communities on 0.5% agarose plates containing M9 minimal media and 0.2% D-glucose to determine the colony forming units (CFU) per ml (fig. S4c). CFU/ml was used as a proxy for total cell number in the community because of the strong correlation with cell counting using a hemocytometer (fig. S4d). The relative contribution of CFU for growth on water alone compared to growth on D-glucose was then used as a relative frequency cutoff for each of the 12 initial communities, respectively (fig. S4e). These values allowed us to estimate lower bounds for community diversity derived from the supplied the carbon source (fig. S6b).

